# The coordination between penicillin-binding protein 1a (PBP1a) and the hydrolytic peptidase DacB determines the integrity of bacterial cell poles

**DOI:** 10.1101/2022.03.18.484884

**Authors:** Huan Zhang, Srutha Venkatesan, Emily Ng, Beiyan Nan

## Abstract

Peptidoglycan (PG) defines cell shape and protects bacteria against osmotic stress. The growth and integrity of PG require coordinated actions between synthases that insert new PG strands and hydrolases that generate openings to allow the insertion. However, the mechanisms of their coordination remain elusive. Here we show that moenomycin that inhibits a family of PG synthases known as Class-A penicillin-binding proteins (aPBPs), triggers cell lysis despite aPBPs being non-essential for cell integrity. We demonstrate that inhibited PBP1a2, an aPBP, accelerates the degradation of cell poles by DacB, a hydrolytic PG peptidase, in the bacterium *Myxococcus xanthus*. Moenomycin reduces the mobility of DacB molecules through PBP1a2, potentially promoting the binding between DacB and PG. Conversely, DacB also regulates the distribution and dynamics of aPBPs. These findings reveal the lethal action of moenomycin and suggest that disrupting the coordination between PG synthases and hydrolases could be more lethal than eliminating individual enzymes.

## Introduction

The peptidoglycan (PG) cell wall, a network of glycan strands and peptide crosslinks, is a hallmark structure of bacteria^1^. PG encases the cytoplasmic membrane, defines cell geometry, and protects the cell against lysis due to its high internal osmotic pressure. As PG is essential for the survival of most bacteria, disrupting PG has become the most successful strategy for antibiotic therapies. The assembly of PG requires multiple synthases, including glycosyltransferases (GTases) that polymerize glycan chains and transpeptidases (TPases) that form peptide crosslinks^2^. During the vegetative growth of most rod-shaped bacteria, PG is assembled by two major enzymatic systems, the Rod system and class A penicillin-binding proteins (aPBPs). The Rod system consists of RodA, a SEDS-family GTase, PBP2, a TPase of the class B penicillin-binding proteins (bPBPs), and MreB, a bacterial actin homolog that orchestrates the actions of RodA and PBP2^3-6^. In contrast, aPBPs possess both GTase and TPase activities and function outside of the Rod system^7,8^. While the Rod system determines rod shape and accounts for the majority of PG growth, aPBPs appear to be secondary and nonessential^3,9,10^. Recent findings suggest that aPBPs could support rod shape indirectly by repairing PG defects and regulating cell diameter^11,12^. However, aPBPs may play more important roles. For instance, the inhibitors of aPBPs usually trigger rapid collapse of rod shape whereas cells can remain rods for generations when the Rod proteins are inhibited or depleted^13-16^.

The insertion of new PG strands requires local hydrolysis of the existing PG network^2,17,18^. PG hydrolases, including lytic transglycosylases, amidases, and endo/carboxypeptidases, are difficult to study in most model organisms. On the one hand, these enzymes are highly redundant in most bacteria, where strains lacking single or several hydrolases usually do not show significant growth or morphological defects. On the other hand, as uncontrolled hydrolytic activities could compromise PG integrity and cause cell lysis, it is difficult to observe highly-activated PG hydrolysis during normal growth^17,19^. To maintain the integrity of cell wall, PG hydrolases must coordinate with synthases in time and space. However, even less is known about the mechanisms of their functional coordination.

*Myxococcus xanthus* is a rod-shaped, gram-negative bacterium that possesses both the Rod system and aPBPs. Similar to other rod-shaped bacteria, the Rod system is essential for the establishment of Rod shape in *M. xanthus*^15,20^. In contrast, aPBPs appear nonessential as the spherical spores of *M. xanthus* can still germinate into rods when aPBPs are inhibited. These newly established rods quickly retrogress to spheres before first division. It remains unknown how aPBPs support rod-like morphology^20^. In this study, we identified DacB, a PBP4-family D-Ala-D-Ala carboxypeptidase/endopeptidase, as a major PG hydrolase that collapses the rod shape of *M. xanthus* by degrading PG, especially at cell poles. We found that whereas either the inhibition or absence of aPBPs is sufficient to enrich DacB to cell poles, only the inhibition, but not absence, of aPBPs triggers the loss of rod shape in vegetative cells. We demonstrate that inhibited PBP1a2, an aPBP, activates PG hydrolysis by DacB by reducing the mobility of DacB molecules. Conversely, DacB also regulates the dynamics and distribution of aPBPs. Our results elucidate the mutual regulation between PG synthases and hydrolases, which plays central roles in the maintenance of cell integrity.

## Results

### The inhibition, but not absence, of aPBPs triggers rapid collapse of rod shape

To study how aPBPs and the Rod system support rod shape, we treated vegetative *M. xanthus* cells using moenomycin (4 μg/ml) and mecillinam (100 μg/ml) at two-fold of their minimum inhibitory concentrations (MICs). Moenomycin is a phosphoglycolipid that inhibits the GTase activity of aPBPs, whereas mecillinam is a ß-lactam that specifically inhibits the Tpase activity of PBP2 in the Rod system^21,22^. After 2 h, 72.7% (n = 534) of moenomycin-treated cells became spherical. In contrast, mecillinam only caused minor bulging near the centers of cells and none (n > 1000) of the treated cells lost rod shape (Fig. 1A). Agents that inhibit PG synthesis or disrupt PG were suspected to induce sporulation of *M. xanthus*^23^. To distinguish whether the spherical cells resulted from moenomycin treatment are spores or vegetative cells that had lost rod shape, we treated these cells with sonication. Compared to the spores from untreated cells where 89 ± 9% (calculated from 3 independent experiments, for each experiment, n > 500, same below) are resistant to sonication^20^, only 5 ± 1% of the moenomycin-induced spherical cells remained after sonication. Thus, our data indicate that the inhibition of aPBPs, but not the Rod system, triggers rapid collapse of rod shape in vegetative *M. xanthus* cells.

**Fig. 1.**
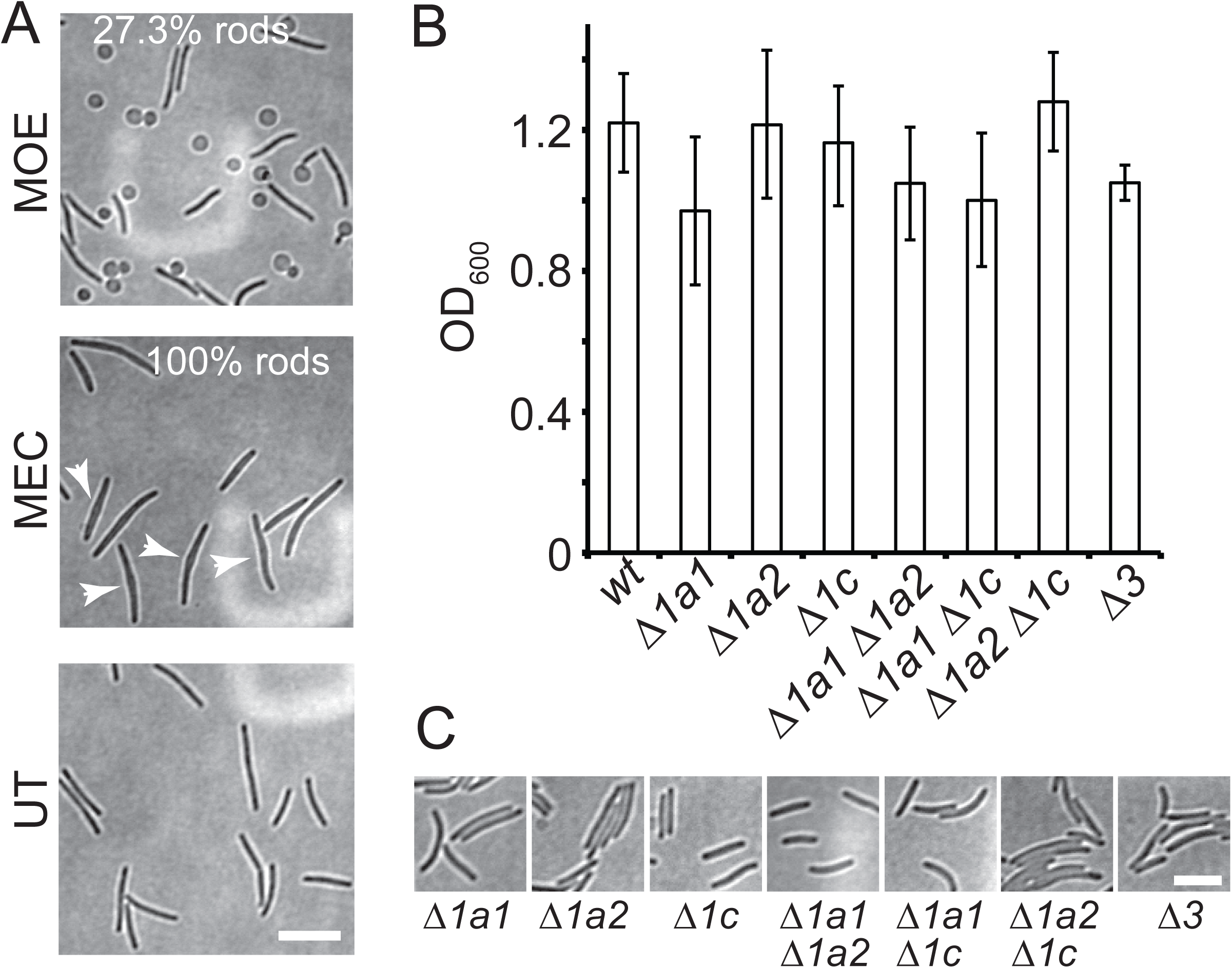
The inhibition, but not absence of aPBPs abolishes rod shape of *M. xanthus*. **A)** Moenomycin (MOE, 4 μg/ml) that inhibits aPBPs induces the loss of rod shape of *M. xanthus*, whereas mecillinam (MEC, 100 μg/ml), an inhibitor of the Rod system, does not. **B)** Cells lacking one, two, or all three (*Δ3*) aPBPs are all viable and only the absence of PBP1a1 causes moderate delay in growth. Cells were inoculated at OD_600_ = 0.2 and growth measured after 12 h of incubation from three biological replicates. **C)**. aPBPs are dispensable for the establishment and maintenance of rod shape. Scale bars, 5 μm.

As moenomycin inhibits aPBPs specifically, we investigated whether the cells that lack certain aPBPs still maintain rod shape. *M. xanthus* encodes three aPBPs, including two homologs of PBP1a (PBP1a1, MXAN_5181 and PBP1a2, MXAN_5911) and one putative PBP1c (MXAN_2419). The strains that lack one, two, or all three aPBPs were all viable and only the absence of PBP1a1 caused moderate delay in growth (Fig. 1B). Importantly, the cells of all seven mutants maintained rod shape (Fig. 1C). Thus, aPBPs are dispensable for both growth and rod-like morphology. It is the inhibition, but not absence of aPBPs that promotes the collapse of rod shape.

### Moenomycin activates PG hydrolysis by DacB

The loss of rod shape in the presence of moenomycin could be a result of blocked PG assembly activities from aPBPs. However, this hypothesis cannot explain the fact that cells completely lacking aPBPs are still rods. Alternatively, the inhibition of aPBPs could activate or alter the distribution of certain hydrolases, which results in misregulated degradation of PG. In a previous attempt to label PG using a fluorescent D-amino acid, TAMRA 3-amino-D-alanine (TADA)^20,24^, we found that deleting *dacB*, a gene encoding a PBP4-family D-Ala-D-Ala carboxypeptidase/endopeptidase, improved the incorporation of TADA significantly^20^ (Fig. 2A, 2B). Thus, consistent with its peptidase activity, DacB actively removes TADA from the peptide stem of PG. To test if DacB participates in the degradation of rod shape in moenomycin-treated cells, we first tested if the *ΔdacB* cells are resistant to moenomycin. We inoculated cells in liquid medium at OD_600_ 0.2 and measured cell concentrations in the presence of moenomycin (2 μg/ml, MIC) after a 24-h incubation. Compared to the wild-type cells that grew very little, the *ΔdacB* strain showed significantly higher resistance (Fig. 2C). We then expressed *dacB* ectopically using a vanilate-inducible promoter^25^ in addition to the endogenous *dacB* gene. Induced by 200 μM vanilate, the cells over-expressing DacB were hyper-sensitive to moenomycin, whereas the ones harboring the empty vector showed sensitivity similar to the wild-type (Fig. 2C).

**Fig. 2.**
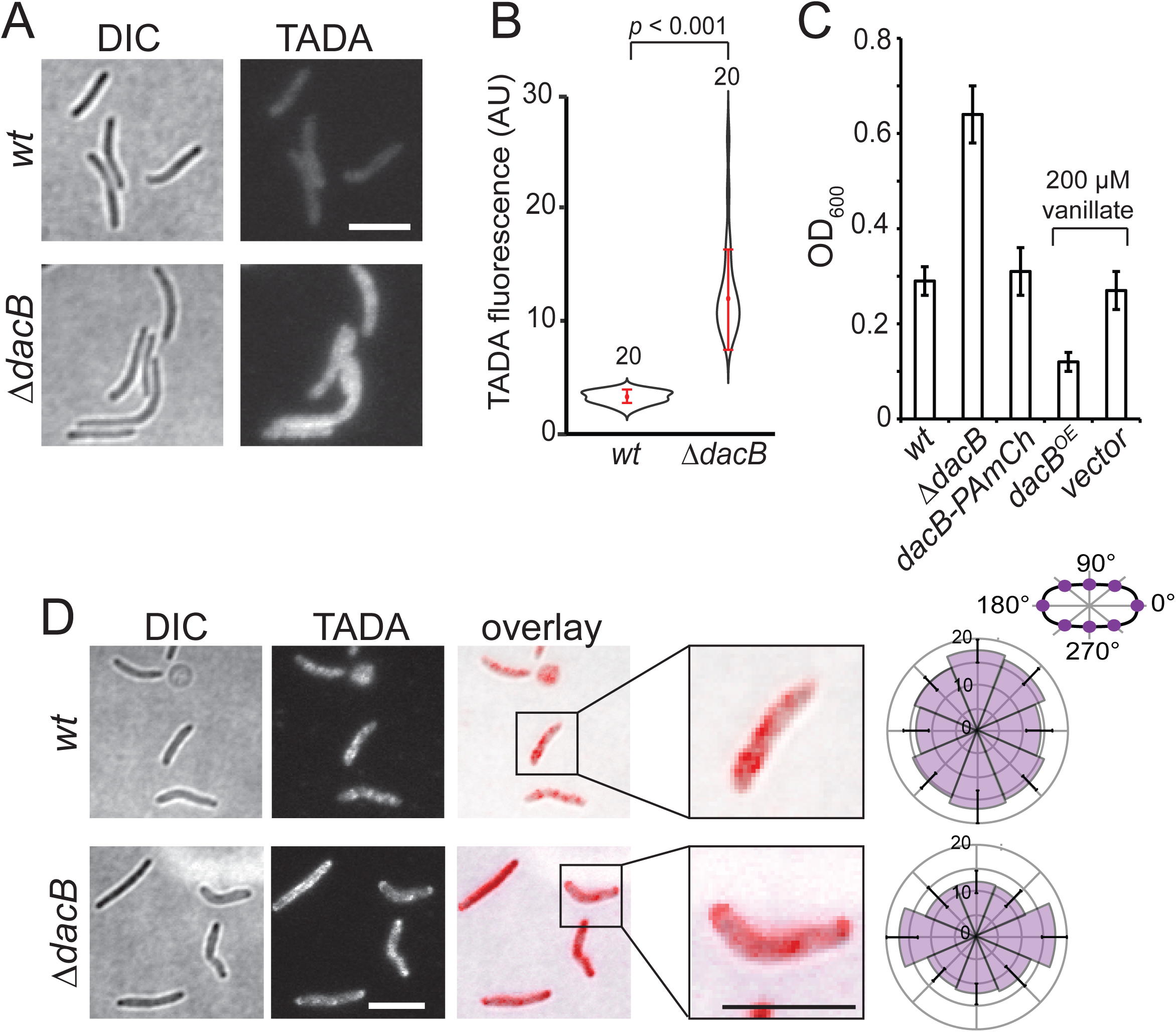
Moenomycin activates PG hydrolysis by DacB, especially at cell poles. **A, B**) DacB is a major PG hydrolase as the absence of DacB enhances the efficiency of TADA incorporation. Vegetative cells of the wild-type and *ΔdacB* strains were imaged using the same setting. The average fluorescence intensity of 20 cells are plotted in **B. C**) Moenomycin activates PG hydrolysis by DacB. The deletion and overexpression of *dacB* enhances and reduces the resistance against moenomycin, respectively. Labeling DacB with PAmCherry does not affect the sensitivity to moenomycin. Vanillate, the inducer for *dacB* overexpression, does not affect the sensitivity as the cells carrying the empty vector show similar sensitivity as the wild type ones. Cells were inoculated at OD_600_ = 0.2 and their concentrations measured from three biological replicates after a 24-h incubation in the presence of moenomycin (2 μg/ml). Whiskers indicate the 25^th^ - 75^th^ percentiles and red dots the median. **D)** Moenomycin induces near even loss of TADA signal in the wild-type cells while the *ΔdacB* cells retain TADA at their poles. The average and standard deviation of TADA intensity were calculated from 20 cells in the diagrams to the right (same below). Scale bars, 5 μm.

To visualize the degradation of PG by DacB, we labeled the entire PG layer of cells by inducing the germination of *M. xanthus* spores in the presence of TADA. Both the wild-type and *ΔdacB* cells showed homogeneous incorporation of TADA (Fig. 2A). After 1 h of moenomycin treatment, the labeled wild-type cells lost fluorescence near evenly throughout the PG layer, whereas bright loci of TADA still remained at the poles in *ΔdacB* cells (Fig. 2D). Taken together, moenomycin activates the hydrolytic activity of DacB, especially at cell poles.

### The inhibition or absence of aPBPs enriches DacB at cell poles

To further investigate the effects of moenomycin on DacB, we fused a DNA sequence encoding photo-activatable mCherry (PAmCherry) to the endogenous gene of *dacB* in wild-type *M. xanthus*. The introduction of PAmCherry did not affect the strain’s sensitivity to moenomycin, indicating that the tagged DacB protein is fully functional (Fig. 2C). We used a 405-nm excitation laser (0.2 kW/cm^2^) to activate the fluorescence of a few labeled DacB molecules randomly in each cell and quantified their localization. Along the long axis of cells, we loosely defined a region within 480 nm from each end of cell as a “pole” and the rest of the cell as the “nonpolar region”. By this definition, poles account for about 20% of the cell surface in a typical *M. xanthus* cell (∼5 μm long, 1 μm wide). We found that in untreated cells, 28.0% (n = 2250) of DacB molecules were detected at cell poles, suggesting that DacB distributes near evenly in the membrane. Consistent with the degradation pattern of PG (Fig. 2C), DacB aggregated at poles (47.7%, n = 3157) in the presence of moenomycin (Fig. 3A). However, despite the fact that cells still retain rod shape (Fig. 1C), we observed similar polar enrichment of DacB in the strains that lacks individual aPBPs (Fig. 1C, 3A). Thus, although the even distribution of DacB requires the presence and activities of all three aPBPs, its polar enrichment is not sufficient to dismantle rod shape.

**Fig. 3.**
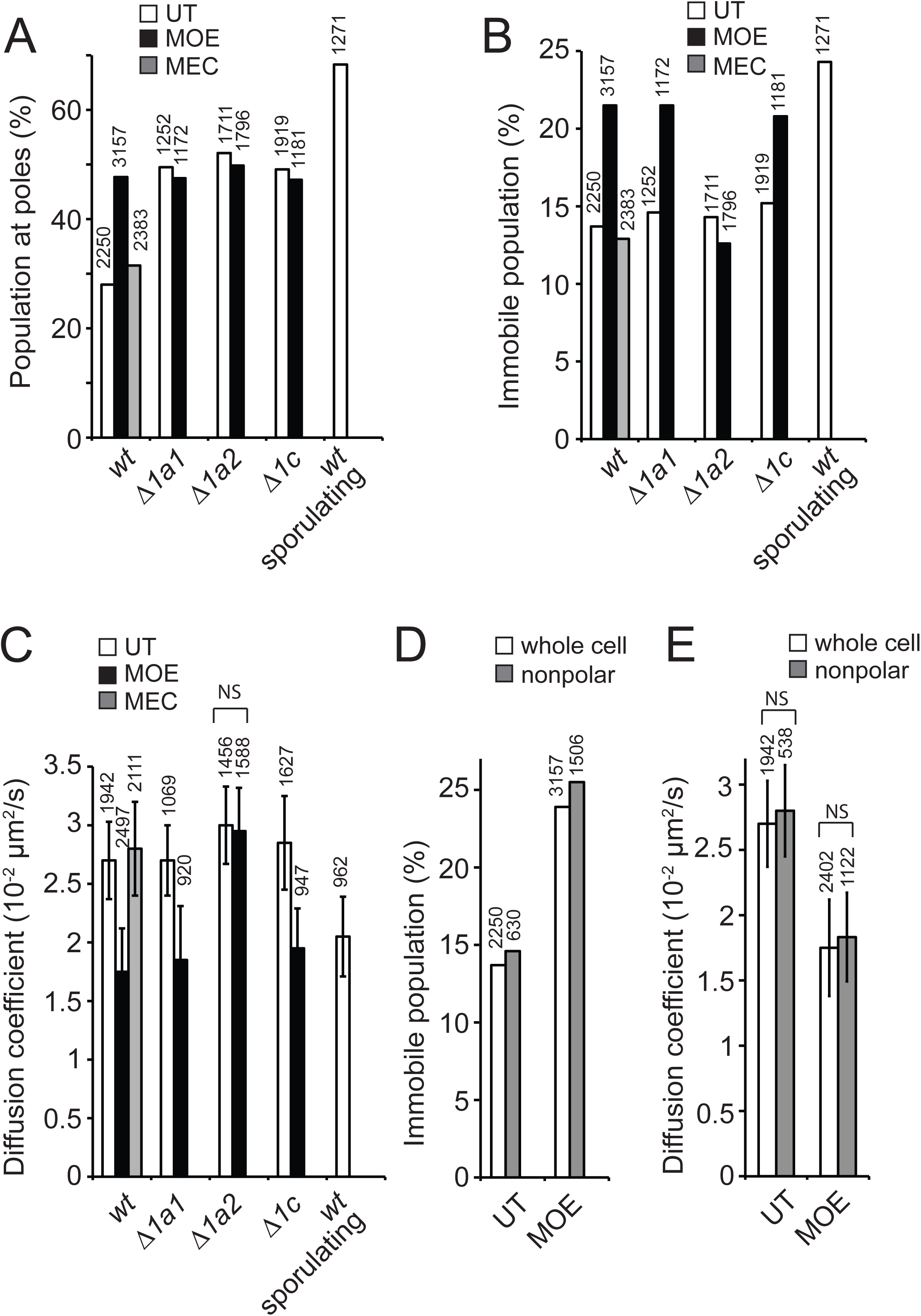
Moenomycin activates DacB through PBP1a2. **A)** The inhibition (by moenomycin (MOE, 4 μg/ml)) and absence of aPBPs enrich DacB to cell poles. **B, C)** Moenomycin increases the population of immobile DacB and decreases the diffusion coefficient of the mobile DacB through PBP1a2. Similar polar enrichment and reduction of mobility of DacB molecules are also observed in the sporulating cells where PG is actively degraded. Mecillinam (MEC, 100 μg/ml) does not affect the distribution and dynamics of DacB (**A-C**). **D, E)** Moenomycin increases the population of immobile DacB (**D**) and reduces the diffusion coefficient of mobile DacB (**E**) throughout the whole cell. The total number of molecules analyzed is shown on top of each bar (same below). NS, nonsignificant difference.

### The mobility of DacB decreases during the degradation of PG

Compared to individual synthases and hydrolases, the existing PG meshwork is a large and stationary substrate. In *Escherichia coli*, individual molecules of aPBPs distribute between immobile and mobile populations and their inhibitors increase their mobility^8^. Similarly, we expect that the mobility of DacB will decrease during its enzymatic action. To test this hypothesis, we imaged single DacB-PAmCherry molecules at 10 Hz using single particle tracking photo-activated localization microscopy (sptPALM)^15,26^. To analyze the data, we only chose the fluorescent particles that remained in focus for 4 - 12 frames (0.4 - 1.2 s). DacB molecules displayed two dynamic patterns, immobile and diffusion. The immobile particles remained within a single pixel (160 nm ×160 nm) before photo-bleach and the mobile ones displayed typical diffusive behavior. In wild-type, untreated vegetative cells, 13.1% (n = 2250) of DacB molecules were immobile and the diffusion coefficient of the mobile population is *D*_*DacB*_ = 2.7 × 10^−2^ ± 4.1 × 10^−3^ μm^2^/s (n = 1955). During moenomycin-induced PG degradation, the immobile population of DacB increased to 20.9% (n = 3157) and the diffusion coefficient of the mobile population decreased to 1.8 × 10^−2^ ± 4.7 × 10^−3^ μm^2^/s (n = 2497). The reduced mobility of DacB suggests an enhanced association between DacB and its PG substrate. As a control, mecillinam that does not trigger immediate PG degradation, did not show significant effects on either the dynamics or distribution of DacB (Fig. 3A-3C).

In response to chemical signals, such as glycerol, vegetative cells of *M. xanthus* thoroughly degrade their PG and transform into spherical spores within two to four hours despite that the expression of the *dacB* gene is not regulated^20,27-30^. If DacB also degrades PG during sporulation, glycerol-induced sporulation could provide a scenario where DacB is highly activated. In fact, using the length/width ratio (L/W) of cells to quantify the sporulation progress, we found that the ***Δ****dacB* cells did not degrade PG efficiently at poles and thus formed spores slowly. By contrast, the over-expression of DacB significantly accelerated the rod-to-sphere transition (Fig. S1). In the sporulating cells where DacB hydrolyzes PG actively, DacB concentrated at poles, its immobile population increased to 23.3% and the diffusion coefficient of the mobile population decreased to 2.1 × 10^−2^ ± 4.4 × 10^−3^ μm^2^/s (n = 1217) (Fig. 3A-3C). As reduced mobility of DacB strongly correlates with enhanced PG degradation in both vegetative and sporulating cells, we reason that the molecular dynamics of DacB could be used to monitor its hydrolytic activity.

### Moenomycin activates DacB through PBP1a2

How does moenomycin activate DacB? To answer this question, we quantified the mobility of DacB in the vegetative cells of the strains lacking individual aPBPs. The absence of each aPBP did not alter either the population of immobile DacB molecules or the diffusion coefficients of the mobile ones (Fig. 3B, 3C). Thus, the absence of each aPBP does not activate DacB. Could moenomycin activate DacB through inhibited aPBPs? If so, DacB will no longer respond to moenomycin in the absence of certain aPBPs. Strikingly, DacB did not respond to moenomycin in the ***Δ****pbp1a2* background, whereas moenomycin still reduced the mobility of DacB in the absence of PBP1a1 and PBP1c (Fig. 3B, 3C). Thus, we conclude that the inhibited PBP1a2 activates PG hydrolysis by DacB.

To determine whether inhibited PBP1a2 activates DacB throughout the whole cell or specifically at cell poles, we studied the dynamics of DacB in nonpolar regions. Similar to its effects on whole cells, moenomycin increased the immobile population to 25.5% (n = 1856) in nonpolar regions and decreased the diffusion coefficient of the mobile population to 1.9 × 10^−2^ ± 3.4 × 10^−3^ μm^2^/s (n = 1383). Thus, inhibited PBP1a2 activates DacB throughout the whole cell and it is the combination of activation and enrichment of DacB that causes the degradation of PG at cell poles (Fig. 3D, 3E).

### DacB regulates the distribution and dynamics of aPBPs

The coordination between PBP1a2 and DacB suggests that DacB could also regulate the activity and distribution of aPBPs. To test this hypothesis, we expressed PAmCherry-labeled aPBPs using their endogenous loci and promoters. All three aPBPs displayed two dynamic patterns, immobile and diffusion (Fig. 4A). Indeed, deletion of the *dacB* gene showed profound effects on the dynamics and spatial distribution of all three aPBPs. The absence of DacB significantly increased and decreased the mobility of PBP1a2 and PBP1c, respectively (Fig. 4A), suggesting that DacB could regulate these aPBPs oppositely. In addition, both PBP1a1 and PBP1c increased their distribution at cell poles in *ΔdacB* cells (Fig. 4B). The regulatory roles of DacB on aPBPs remain to be investigated.

**Fig. 4.**
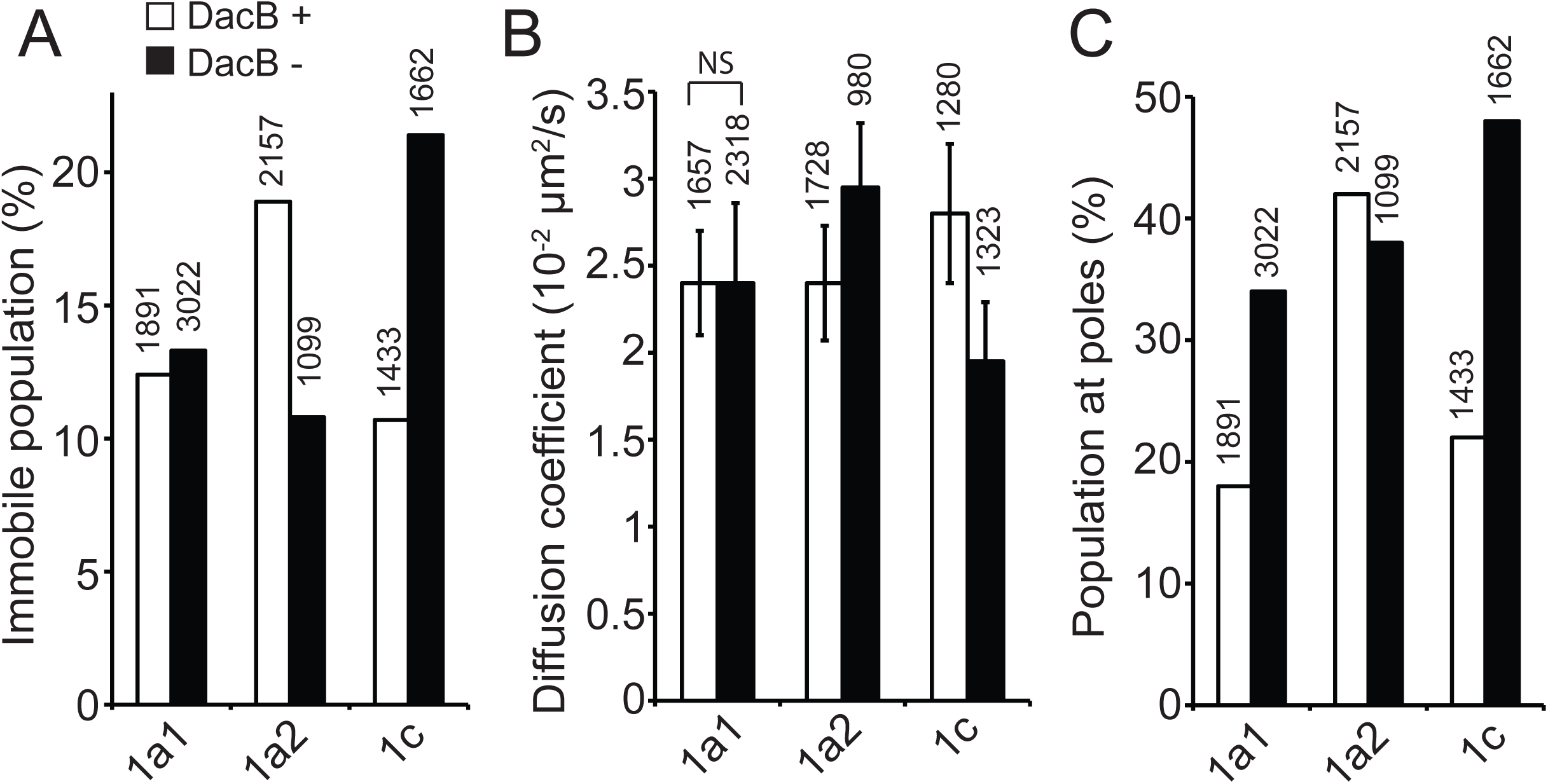
DacB regulates the dynamics and distribution of aPBPs. **A)** The absence of DacB decreases and increases the immobile populations of PBP1a2 and PBP1c, respectively. **B)** The absence of DacB increases and decreases the diffusion coefficient of the mobile molecules of PBP1a2 and PBP1c, respectively. **C)** The absence of DacB enriches PBP1a1 and PBP1c at cell poles. NS, nonsignificant difference.

## Discussion

### A quantitative method to study the coordination between PG synthases and hydrolases

It has long been speculated that PG hydrolases associate with synthases in many functions that maintain the integrity of PG^17,19,31^. Hydrolases could even determine the localization and activities of synthases^31^. Although nearly all the PG synthases and hydrolases have been identified, their functional coordination is largely unknown. The major hurdle in such research is that aPBPs and PG hydrolases are highly redundant and many single mutations do not show clear phenotypes. Furthermore, instead of forming stable complexes, these enzymes that carry opposite activities could interact transiently. In *E. coli*, some endopeptidases and a lytic transglycosylase interact with PBP1a and PBP1b through protein adaptors^32,33^. However, the search for such adaptors failed to yield positive results in *M. xanthus*. Thus, although the coordination between PG synthases and hydrolases could be universal, its mechanisms could vary in different organisms. In addition, we found that cells lacking aPBPs and DacB are viable, whereas the changes in their interaction cause cell lysis. For these reasons, genetic and biochemical approaches alone might not be sufficient for studying the complex coordination among these enzymes. This research has provided a method to study such coordination before the identification of the coupling mechanisms. We quantified the localization and dynamics of single enzymes and presented evidence for the mutual regulation between an aPBP and a PG hydrolase. Future work will be done to investigate whether or not the Rod system coordinates with specific PG hydrolases.

### The roles of aPBPs and cell poles in rod-like morphology

Recent researches on rod-like morphology has emphasized the roles of the Rod system as it is the major machinery that assembles PG on the cylindrical surface of the cell^2,18,34^. In contrast, cell poles are considered inert after division and their roles in the integrity of cell wall are less appreciated^35^. Here we show that poles are critical for rod-like morphology, as PG hydrolysis at poles causes rapid collapse of rod shape. Our findings also provide an explanation for the lytic effects of aPBP inhibitors. The absence of the Rod system from cell poles, probably due to the exclusion of MreB^36^, makes aPBPs the dominant PG synthases at polar regions. As a consequence, when aPBPs are inhibited, poles become void of PG synthases. The lack of PG repair in combination with the enrichment of activated DacB by PBP1a2, causes rapid PG degradation at poles, which results in sudden loss of rod shape. In contrast, the cylindrical surfaces of cells are maintained by both the Rod system and aPBPs. Thus, when one system is inhibited, the other is still able to repair PG damages^12^.

### The bactericidal action of moenomycin

Despite exhaustive data on how antibiotics exert action on their targets, we know very little about how these drugs actually kill bacteria^37^. For instance, ß-lactams specifically inhibit the TPase activity of PBPs. However, rather than the blockage of PG assembly, the actual lethal effect of ß-lactams is inducing the degradation of uncrosslinked glycan strands, which are produced by the uninhibited GTases^38^. Similarly, blocking the GTase activities of aPBPs is not likely the direct bactericidal effect of moenomycin. Similar to *M. xanthus*, although wild-type *Bacillus subtilis* is sensitive to moenomycin, a strain lacking all four aPBPs is still viable and produces PG with only minor defects^3,10^. In this study, we provide evidence that moenomycin inhibits PBP1a2, which in turn, regulates the activity and distribution of DacB and eventually causes cell lysis. We propose that the bactericidal action of moenomycin is inducing an imbalance between aPBPs and PG hydrolases. Our results suggest that disrupting the coordination between PG synthases and hydrolases could be an effective strategy for antibacterial therapies.

## Materials and Methods

### Strains and growth conditions

Vegetative *M. xanthus* cells were grown in liquid CYE medium (10 mM MOPS pH 7.6, 1% (w/v) Bacto™ casitone (BD Biosciences), 0.5% yeast extract and 8 mM MgSO_4_) at 32 °C, in 125-ml flasks with rigorous shaking, or on CYE plates that contains 1.5% agar. Deletion and insertion mutants were constructed by electroporating *M. xanthus* cells with 4 μg of plasmid DNA. Transformed cells were plated on CYE plates supplemented with 100 μg/ml sodium kanamycin sulfate or 10 μg/ml tetracycline hydrochloride. All constructs were confirmed by PCR and DNA sequencing.

### Sporulation and TADA-labeling

To induce sporulation, glycerol was added to 1 M when liquid cell culture reaches OD_600_ 0.1 – 0.2. After rigorous shaking overnight at 32 °C, remaining vegetative cells were eliminated by sonication and spores were purified by centrifugation (1 min, 15,000 g and 4 °C). The pellet was washed three times with water. To prepare TADA-labeled vegetative cells, purified spores were diluted to OD_600_ 0.5 into 1 ml of liquid germination CYE (CYE medium supplemented with 150 μM TADA, additional 0.2% casitone and 1 mM CaCl_2_) and incubated in an 18-mm test tube at 32 °C, with vigorous shaking.

### Microscopy Analysis

For all imaging experiments, 5 μl cells were spotted on an agarose pad. For the treatments with inhibitors, inhibitors were added into both the cell suspension and agarose pads. The length and width of cells were determined from differential interference contrast (DIC) images as described^20^. DIC and fluorescence images of cells were captured using a Hamamatsu ImagEM X2™ EM-CCD camera C9100-23B (effective pixel size 160 nm) on an inverted Nikon Eclipse-Ti™ microscope with a 100× 1.49 NA TIRF objective. For sptPALM, *M. xanthus* cells were grown in CYE to 4 ×10^8^ cfu/ml. PAmCherry was activated using a 405-nm laser (0.3-3 W/cm^2^), excited and imaged using a 561-nm laser (0.2 kW/cm^2^). Images were acquired at 10 Hz. Single PAmCherry particles were localized, fit by a symmetric 2D Gaussian function, and their trajectories were analyzed as previously described^15^.

## Acknowledgements

We thank Dr. Sofiene Seef and Tâm Mignot for the ***Δ****dacB* strain, Drs. Michael Van Nieuwenhze and Yen-Pang Hsu for providing TADA, and Joshua Pettibon for critical reading of this manuscript. This work is supported by the National Institute of Health R01GM129000.

## Figure Legends

**Fig. S1. DacB is a major PG hydrolase that dismantles rod shape during the sporulation of *M. xanthus*. A)** The absence and overexpression of DacB delays and accelerates the sporulation of *M. xanthus*, respectively. Phase contrast images of cells after 2-h of glycerol induction are shown. **B)** The progress of sporulation is quantified by the length/width ratio of cells after 2-h of glycerol induction. Whiskers indicate the 25^th^ - 75^th^ percentiles and red dots the median. The total number of cells analyzed is shown on top of each bar. **C)** While wild-type cells lose TADA signal near evenly during sporulation, the *ΔdacB* cells retain TADA at their poles. The average and standard deviation of TADA intensity were calculated from 20 cells in the diagrams to the right. Scale bars, 5 μm.

