## Supplementary figures and images for "The coordination between penicillin-binding protein 1a (PBP1a) and the hydrolytic peptidase DacB determines the integrity of bacterial cell poles"

### Fig. S1

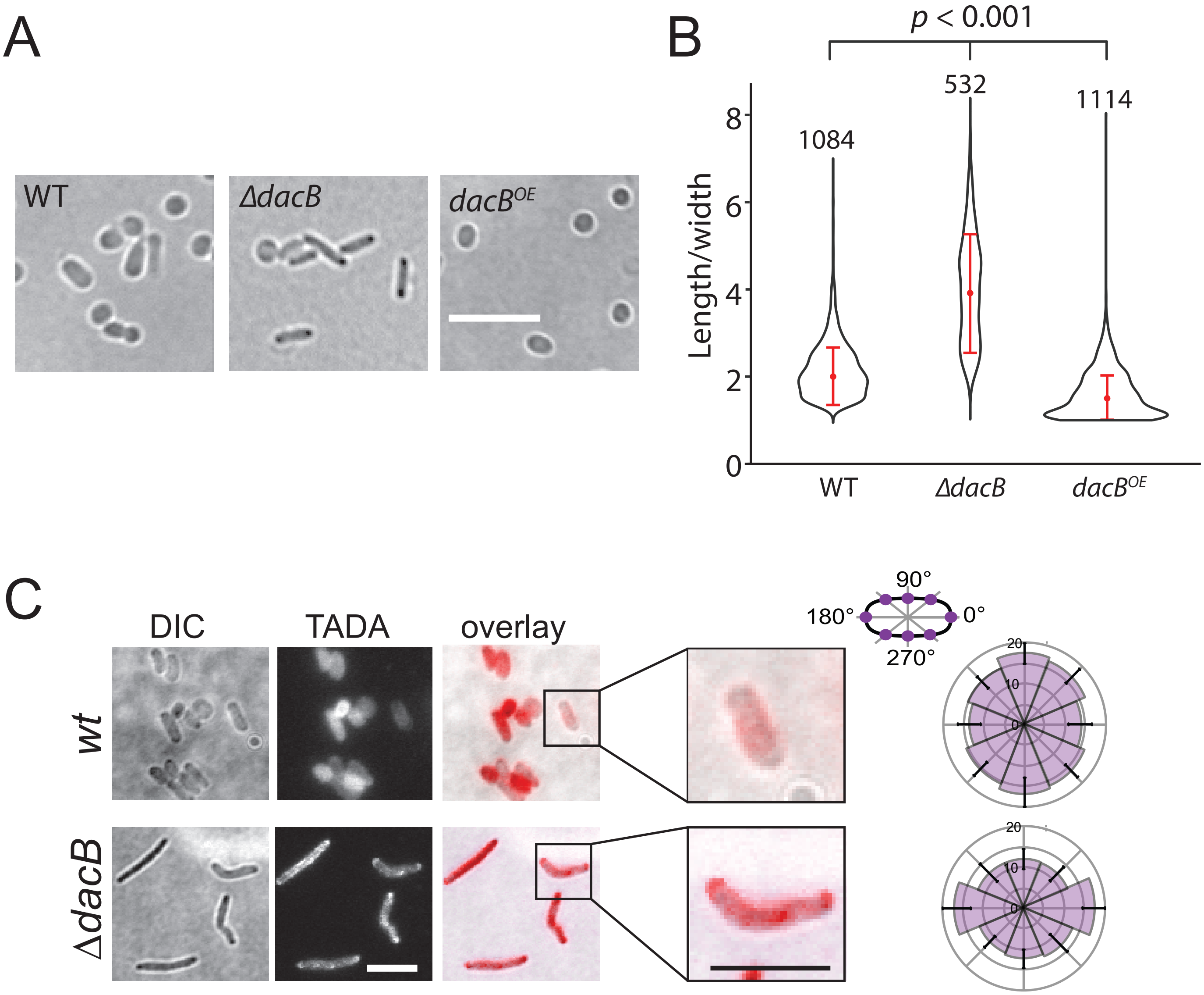
